# Spatiotemporal dynamics of tumor - CAR T-cell interaction following local administration in solid cancers

**DOI:** 10.1101/2024.08.29.610392

**Authors:** Katherine Owens, Aminur Rahman, Ivana Bozic

## Abstract

The success of chimeric antigen receptor (CAR) T-cell therapy in treating hematologic malignancies has generated widespread interest in translating this technology to solid cancers. However, issues like tumor infiltration, the immunosuppressive tumor microenvironment, and tumor heterogeneity limit its efficacy in the solid tumor setting. Recent experimental and clinical studies propose local administration directly into the tumor or at the tumor site to increase CAR T-cell infiltration and improve treatment outcomes. Characteristics of the types of solid tumors that may be the most receptive to this treatment approach remain unclear. In this work, we develop a spatiotemporal model for CAR T-cell treatment of solid tumors, and use numerical simulations to compare the effect of introducing CAR T cells via intratumoral injection versus intracavitary administration in diverse cancer types. We demonstrate that the model can recapitulate tumor and CAR T-cell data from imaging studies of local administration of CAR T cells in mouse models. Our results suggest that locally administered CAR T cells will be most successful against slowly proliferating, highly diffusive tumors, which have the lowest average tumor cell density. These findings affirm the clinical observation that CAR T cells will not perform equally across different types of solid tumors, and suggest that measuring tumor density may be helpful when considering the feasibility of CAR T-cell therapy and planning dosages for a particular patient. We additionally find that local delivery of CAR T cells can result in deep tumor responses, provided that the initial CAR T-cell dose does not contain a significant fraction of exhausted cells.

## 1. Introduction

CAR T-cell therapy involves genetically modifying a patient’s T cells to better detect and mount an attack against cancer cells. This technology has demonstrated unprecedented success in combating previously untreatable hematologic malignancies, with response rates of 40-98% [1], and six drugs have been approved by the Federal Drug Administration since 2017 [1]. The success of CAR T-cell therapies against blood cancers has generated significant interest in adapting CAR T-cell technology to treat solid tumors [2, 3]. Advances in CAR T-cell specificity [4], endurance [5, 6], and trafficking [7, 8] are being investigated in order to overcome the challenges presented by the complex biology of solid tumors. A growing body of work has also focused on local delivery of CAR T cells, which can allow treatment to bypass physical barriers, shorten the distance travelled by T cells, and reduce on-target/off-tumor effects[9–12]. This could simultaneously increase the number of CAR T cells within the tumor and decrease the risk of immune-related toxicity. Expanding evidence from pre-clinical and phase I clinical trials also suggests that local delivery of CAR T cells is safe and feasible[13–20].

The efficacy of intracranial or intratumoral delivery of CAR T cells has been demonstrated in several mouse models of brain tumors [9–11, 13], and in case studies in human patients [21–23]. Further studies in mice have tested regionally delivered CAR T cells against breast cancer and liver cancer [12, 24], and in some cases demonstrated that regional delivery reduces the required CAR T cell dose for effective therapy compared with systemic administration [25, 26]. A number of ongoing clinical trials utilize regional delivery of CAR T cells, with the majority targeting tumors of the central nervous system or liver [16–20, 27**?**]. Additional locoregional delivery trials target malignant pleural mesothelioma [15, 28, 29], head and neck cancers [30], and breast cancer [31].

Mathematical models of CAR T-cell treatment have enabled characterization of CAR T cell population dynamics [32, 33], CAR T-cell proliferation and/or killing mechanisms [34–39], and have been used to predict patient outcomes [40–43], and suggest optimal treatment plans [42, 44–49]. Tserunyan et al. and Nukula et al. provided comprehensive reviews on the subject [50, 51]. Most existing models yield helpful insight under the assumption that cell populations are well-mixed throughout the course of treatment. However, this assumption is unlikely to hold during CAR T-cell treatment of solid tumors, where failure of CAR T cells to infiltrate the tumor can be a mode of tumor escape [7]. Spatial variability is particularly relevant in the case of regional delivery of CAR T cells, where they may be injected directly into the center of the tumor or into a cavity containing the tumor. To account for spatial variation in cell concentration and provide insight into how the geometry of the delivery of CAR T cells impacts treatment efficacy, we employ a 3-D reaction-diffusion modeling framework for local administration of CAR T cells to treat solid tumors.

Reaction-diffusion equations have been used to model tumor growth for nearly three decades. Murray et al. pioneered this approach [52], writing out in words that

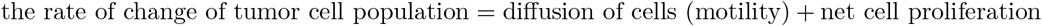

This formulation inspired a rich body of work, particularly successful in modelling highly diffusive cancers like glioma, and their treatments including resection and chemotherapy [53–56]. A similar modeling paradigm was proposed by Matzavinos et al. to study the immune response to a dormant tumor [57] and Li et al.[58] to study the infiltration of T cells into breast cancer tumor cell clusters. Reaction-diffusion equations have further been used to model nanoparticle drug delivery and response [59–61], and to model intratumoral injection of ethanol [62, 63]. However, the post-injection kinetics of adoptive cellular therapies like CAR T cells are fundamentally different than those of purely diffusive, small-molecule based drugs. As a “living drug,” infused cells will ideally proliferate and persist in the body in a way that traditional pharmacokinetic modeling frameworks do not capture.

In this article, we develop a 3-dimensional spatio-temporal model for CAR T-cell therapies applied to dynamic solid tumors. We compare model simulations with experimental data from CAR T-cell imaging studies in mice to verify that model behavior is reasonable. We use this framework to compare two modes of locoregional delivery, intracavitary injection versus intratumoral injection. In particular, we discuss how tumor characteristics like growth rate and invasiveness impact the effectiveness of CAR T-cell treatment.

## 2 Results

### 2.1 Model Overview

We developed a partial differential equations (PDE) model with thresholded diffusion, non-linear growth, and non-linear coupling to describe the spatial coevolution of tumor and CAR T cells in time (See schematic in Fig.1A and details in *Materials and Methods*). In our model, tumor cells both proliferate and diffuse in a density-dependent manner to reflect two stages of tumor growth: an initial phase in which the tumor grows non-diffusively, followed by a diffusive growth phase [55]. We model tumor growth logistically inducing exponential proliferation at low tumor densities and transitioning to a decreased rate as the tumor density at a given point in the spatial domain approaches a local carrying capacity of 1*/b*. When the tumor density exceeds a tumor diffusion threshold, *u*^*^, tumor cells start to diffuse away from that spatial location.

CAR T cells can kill tumor cells upon interaction, proliferate when tumor cell lysis is occuring, become exhausted through interaction with tumor cells, and die. Their movement is described by simple diffusion [64]. Assuming spherical symmetry greatly reduces the complexity of the model, while still allowing us to test two medically relevant methods of local CAR T-cell administration: intracavitary injection (uniform concentration along the boundary of the tumor) and intratumoral injection (high concentration in the center of the tumor). This geometry could also be used to consider treatment plans involving multiple doses, combinations of intratumoral and intracavitary administration, and CAR T-cell injection following tumor resection. Illustrations of the initial CAR T-cell profile for the two administration methods that we will compare are given in Fig. 1B-C. In order to carry out informative numerical analysis and simulations of the model, we estimated several parameter values from experimental data and identified others from previous modeling work (Materials and Methods).

**Figure 1:**
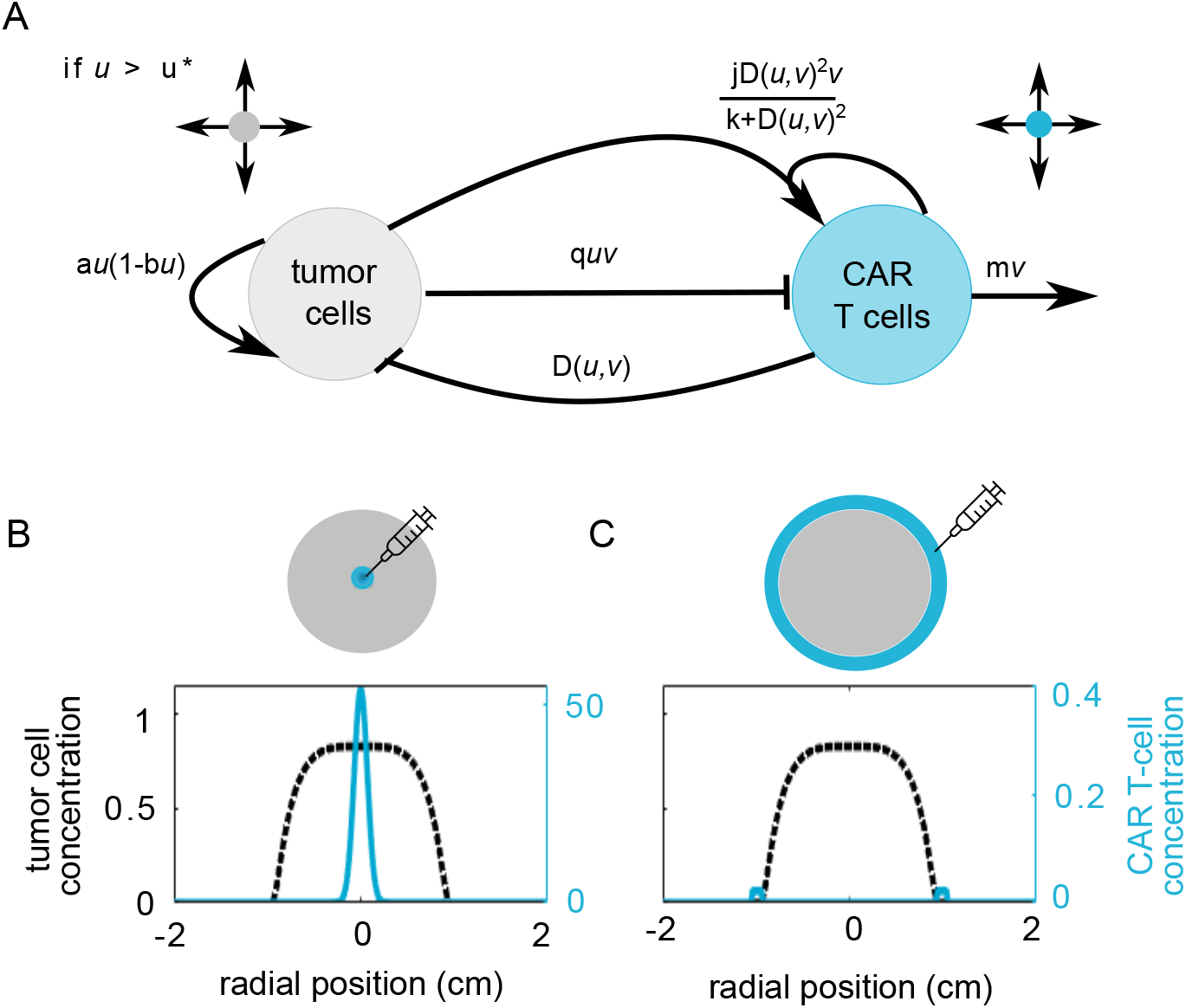
(A) Schematic of PDE model for CAR T-cell treatment of solid tumors. Tumor cells grow logistically and diffuse if their density is above threshold *u*^*^. CAR T cells colocalized with tumor cells can kill tumor cells at lysis rate *D*(*u, v*), but also become exhausted through interacting with tumor cells at rate *quv*. CAR T cell proliferation is driven by tumor cell lysis, and saturates at rate *jv*. CAR T cells disperse via simple diffusion and also senesce at rate *mv*. We model two possible methods of local administration: (B) intratumoral injection, initiated with a high concentration of CAR T cells at the center of the tumor mass and (C) intracavitary injection, initiated with a layer of CAR T cells on the surface of the tumor mass.

### 2.2 Tumor growth in the absence of treatment

Our model provides a flexible framework to simulate a range of solid tumor behaviors. Here we consider four tumor types defined by a unique combination of a proliferation rate, *a*, diffusion coefficient, 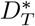, and diffusion threshold, *u*^*^ (See Table S1). For each tumor type, we generated a tumor density profile to represent the tumor burden at the time of CAR T-cell treatment. We initiated the simulation with a spherical tumor of radius 1 mm and uniform density at the carrying capacity, 1*/b*, and then ran it forward in time until the detectable tumor radius reached 1 cm (Fig. 2A). From these initial conditions, we simulated the progression of the tumor over the course of 200 days in the absence of treatment (Fig. 2B). Tumor type I has a high proliferation rate and lower diffusivity resulting in a tumor with a volume doubling time (VDT) of 104 days and an almost uniform, highly dense profile. This type of growth pattern represents compact solid tumors, which could describe some liver or breast cancers [65, 66]. Tumor type II has a moderate proliferation rate and moderate diffusivity, which results in a shorter VDT of 63 days and a slightly less compact cell density profile. This class could represent more aggressive forms of breast cancer [66]. Tumor type III has a low proliferation rate but high diffusivity, resulting in a VDT of 33 days. This balance of proliferation and diffusion generates an invasive but low density growth front that is able to advance undetected, behavior characteristic of a subset of gliomas [67]. Finally, tumor type IV is the most aggressive with both high proliferation and a high diffusivity. More advanced, aggressive gliomas may follow this growth pattern, which has a VDT of only 17 days [67].

**Figure 2:**
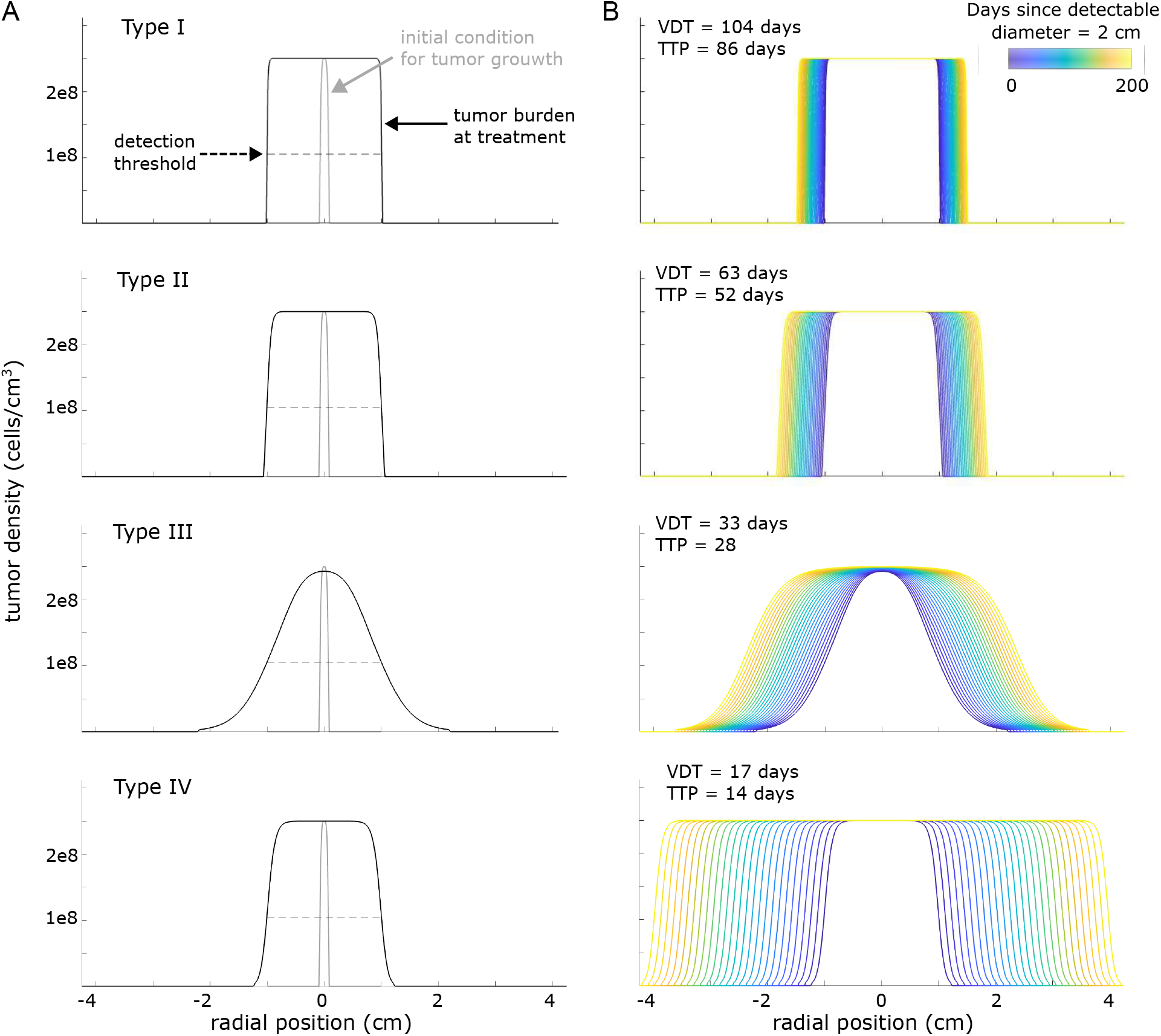
Tumor growth in the absence of treatment for 4 tumor types. Panel (A) shows the density profile for each of these tumors when initiated from a 1 mm mass (gray line) and allowed to grow until the detectable diameter reaches 2 cm (black line) when using a detection threshold around 10^8^ cells/cm^3^. If these four tumors remain untreated, their expansion would continue. In panel (B) we illustrate the growth of the untreated tumors over 200 days. The volume doubling time (VDT) and time to progression (TTP) were calculated from the tumor with detectable diameter of 2 cm as a baseline. Type I proliferates rapidly and diffuses slowly (0.25 day^−1^, 1*/b* = 2.39e8 cells/cm^3^, 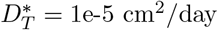, *u*^*^ = 1.2e8 cells/cm^3^) Type II both proliferates and diffuses at a moderate rate (*a* = 0.125 day^−1^, 2.39e8 cells/cm^3^,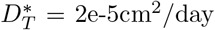, *u*^*^ = 2.39e7 cells/cm^3^), Type III proliferates slowly but diffuses rapidly (*a* = 0.025 day^−1^, 1*/b* = 2.39e8 cells/cm^3^, 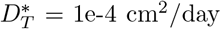, *u*^*^ = 2.39e6cells/cm^3^), and Type IV both proliferates and diffuses rapidly (*a* = 0.25 day^−1^, 1*/b* = 2.39e8 cells/cm^3^, 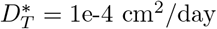, *u*^*^ = 2.39e6cells/cm^3^).

We assessed the impact of varying the tumor proliferation rate, tumor diffusion constant, and tumor diffusion threshold on VDT and the tumor burden at detection. VDT is more sensitive to the tumor diffusion constant than to the tumor proliferation rate (top row, Fig. S1). For tumors with a low proliferation rate and moderate to high diffusivity the tumor diffusion threshold modulates the extent of the tumor that is undetectable (bottom row, Fig.S1 A), whereas for tumors with higher proliferation rates the tumor burden at detection remains relatively constant despite changing the diffusion threshold (bottom row, Fig. S1 B-C).

### 2.3 Intratumoral injection can lead to tumor eradication

We simulated intratumoral CAR T-cell injection by starting with a tumor of diameter 2 cm, constructed using growth parameters from one of the four tumor types defined in the previous section, and a high concentration of CAR T cells in the center of the tumor (see Materials and Methods for details). We observe that CAR T cells injected intratumorally rapidly kill cells at the center of the tumor and start to diffuse out towards the surface, both proliferating and killing tumor cells as they travel (Fig. 3 A). The outcome of treatment strongly depends on whether CAR T cells can proliferate and diffuse sufficiently to reach the surface of the tumor before becoming exhausted. In numerical simulations, we observe four potential treatment outcomes, defined by clinical criteria of response in solid tumors[68]. At sufficiently large doses of effector CAR T cells, intratumoral injection eliminates the tumor, leading to complete response (Fig. 3A-B). For tumor types I-III a partial response is possible in which the CAR T cells reduce the detectable tumor radius significantly, but relapse occurs rapidly after effector CAR T cells dissipate allowing tumor cells at the edge of the initial tumor to proliferate (Fig. 3C-D). Such rapid relapse following CAR T-cell treatment has been observed in the clinic for hematological malignancies. For example, 49% percent of relapses after CAR T-cell treatment in relapsed/refractory DLBCL occur within the first month [69]. With lower doses, CAR T cells may proliferate transiently but allow tumor escape. In these cases, CAR T cells reduce tumor density at the core, but the detectable tumor radius remains unaffected because the T cells do not reach the surface of the tumor in sufficient numbers to contain tumor growth (Fig. 3E-F). With more aggressive tumors, disease progression following failed treatment occurs even faster. For even lower doses of CAR T cells, the injected CAR T cells barely proliferate, instead dying off with little to no noticeable impact on the tumor. Failure of CAR T cells to successfully engraft and proliferate is also a documented failure mode, particularly in the hostile solid tumor micro-environment [70].

**Figure 3:**
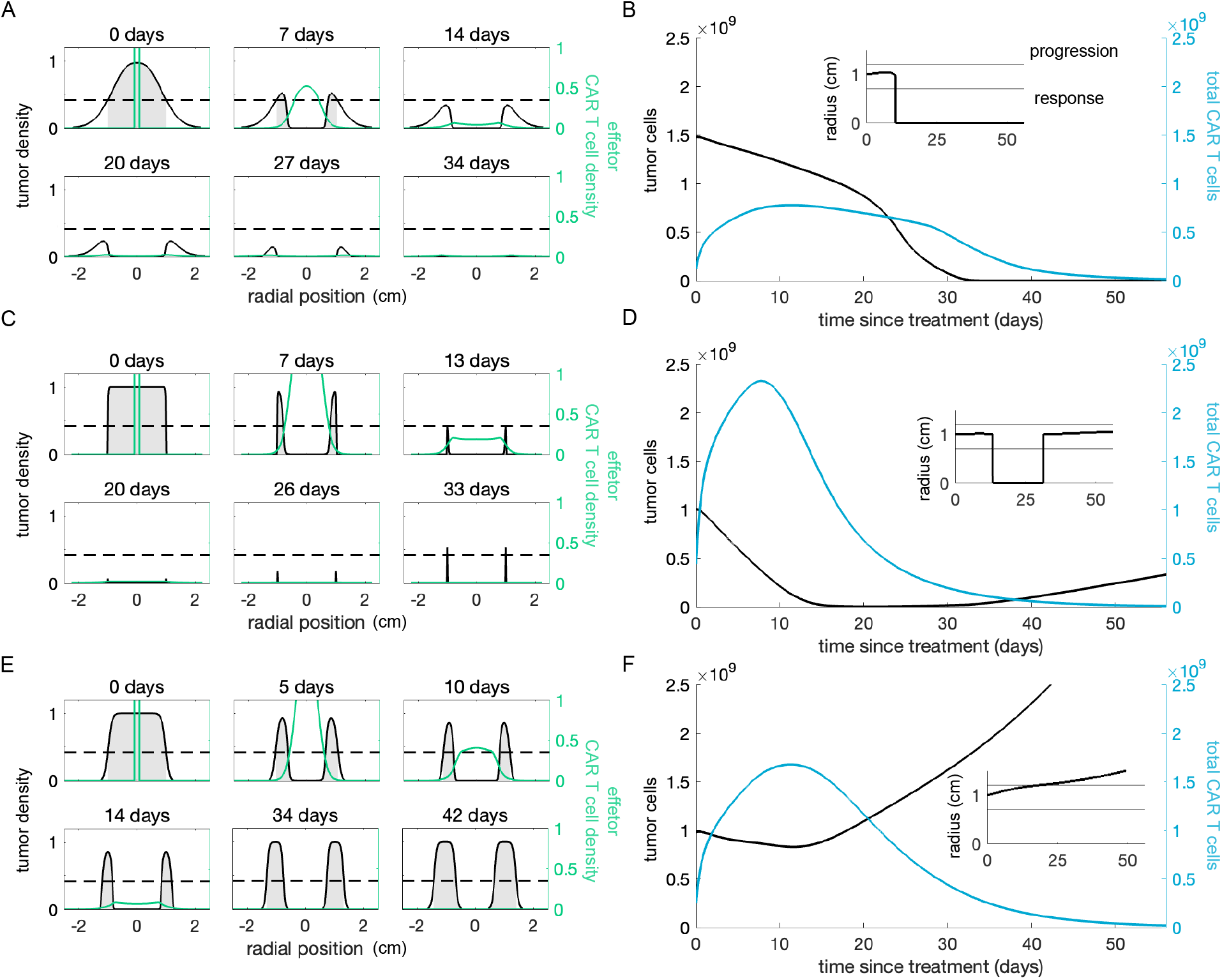
Characteristic intracavitary treatment outcomes. The first column shows snapshots of the cell density profiles following CAR T cell injection. For each of these examples we assumed that 100% of the injected CAR T cells are effectors, and only the density of effector CAR T cells is depicted. The tumor detection threshold is indicated with a dotted line and the portion of the tumor that is detectable is shaded in gray. The second column shows the tumor and total CAR T cell population over time with the corresponding detectable tumor radius inset. (A-B) Treating tumor type I with an intratumoral dose of 1 × 10^9^ CAR T cells reduces the detectable tumor burden and total tumor cell count; however, the tumor resumes growing once CAR T cells dissipate. (C-D) Intratumoral injection of 1 × 10^9^ CAR T cells to treat tumor type III holds the tumor cell count and detectable diameter steady over 8 weeks following treatment. (E-F) Treating tumor type IV with 2 × 10^8^ cells, kills tumor cells at the edge of the initial mass, but the tumor continues to grow uncontrolled

### 2.4 Intracavitary delivery can delay tumor progression

Intracavitary administration of CAR T cells involves delivering CAR T cells into the region surrounding a tumor, but not directly into the tumor itself. For example, in one patient with recurrent glioblastoma, CAR T cells were infused intracranially directly through a catheter device [21]. Another example is delivering CAR T cells via hepatic artery infusion to treat tumors in the liver [27]. In contrast with intratumoral injection, in our model simulations CAR T cells delivered intracavitarily contain the tumor at the growth front as they migrate inwards across the tumor from the outer edge towards the center. To simulate intracavitary delivery of CAR T cells, we initiate the system with one of the four final tumor profiles in Fig. 2A, and a thin layer of CAR T cells concentrated on the surface of the tumor. In our simulations using realistic clinical doses, CAR T cells delivered intracavitarily never eradicate tumors with an initial diameter of 2 cm or more. However, sufficiently large doses can reduce the detectable tumor radius of tumor types I and II, and consequently delay the time to tumor progression. An example of one such scenario is depicted in Figure 4A-B. Against tumor types II and III, moderate to large doses of CAR T cells delivered intracavitarily can hold the tumor to a steady size for an extended period of time. For tumor type III, large doses markedly change the density profile of the tumor, making it more compact (Fig. 4C-D). At low doses against all tumors, or even moderate doses against the most aggressive tumors, eg. type IV, intracavitary delivery of CAR T cells has a minimal impact on tumor progression (Fig. 4E-F).

**Figure 4:**
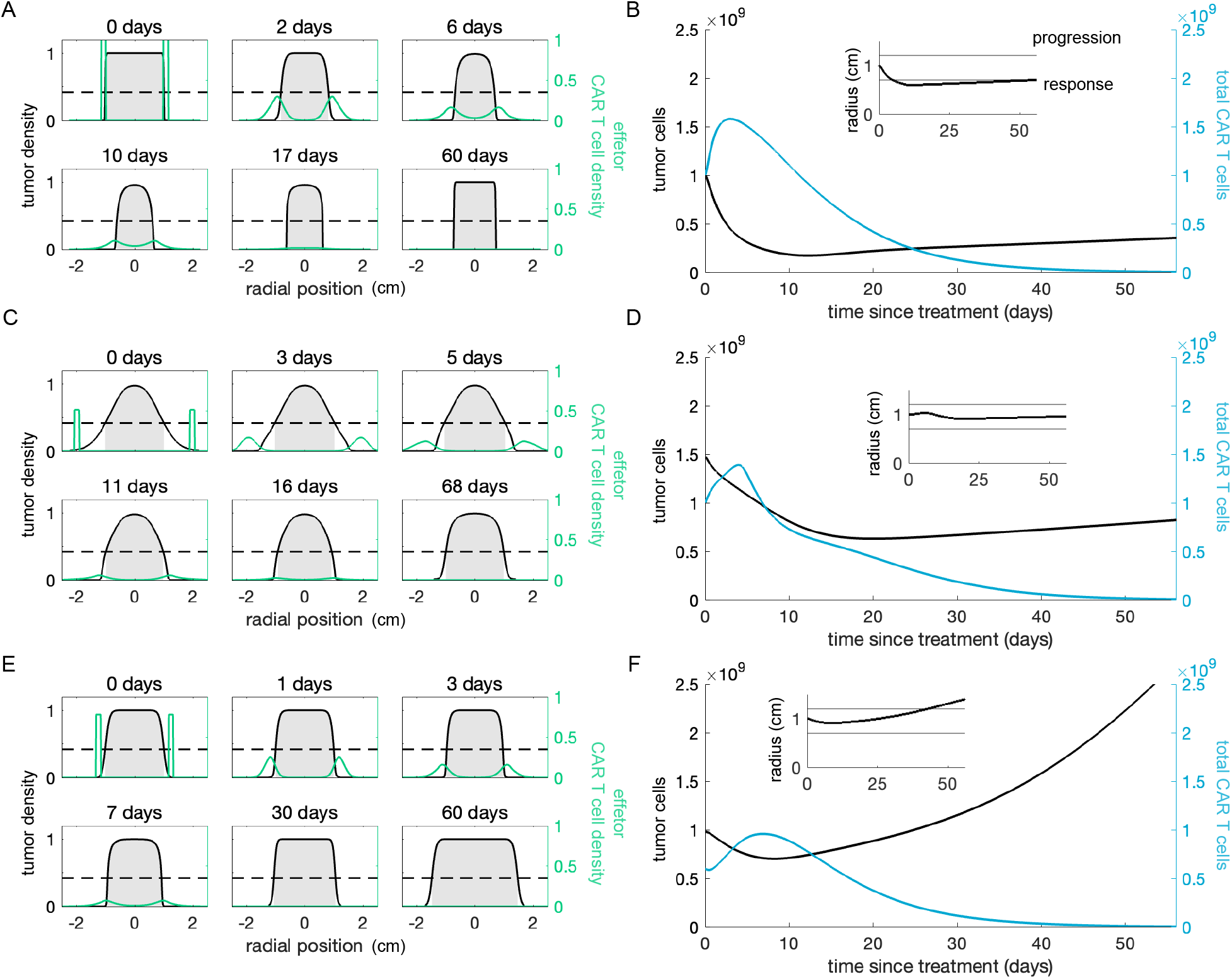
Characteristic intracavitary treatment outcomes. The first column shows the tumor and total CAR T cell population over time with the corresponding detectable tumor radius inset. For each of these examples we assumed that 100% of the injected CAR T cells are effectors. The second column shows snapshots of the cell density profiles over time. Only the density of effector CAR T cells is depicted. The detection threshold is indicated with a dotted line and the portion of the tumor that is detectable is shaded in gray. (A-B) Treating tumor type I with an intratumoral dose of 1 × 10^9^ CAR T cells reduces the detectable tumor burden and total tumor cell count; however, the tumor resumes growing once CAR T cells dissipate. (C-D) Intratumoral injection of 1 × 10^9^ CAR T cells to treat tumor type III holds the tumor cell count and detectable diameter steady over 8 weeks following treatment. (E-F) Treating tumor type IV with 2 × 10^8^ cells, kills tumor cells at the edge of the initial mass, but the tumor continues to grow uncontrolled

### 2.5 Model recapitulates data from experiments in mice

Having established the general behavior of our model, we next verified that it could recapitulate data from two studies in mice: one testing the efficacy of CAR T cells injected directly into solid tumors[71] and another comparing the efficacy of regional vs. systemic delivery of CAR T cells in treating solid tumors[72]. Zhao et al. established large flank mesothelioma tumors in mice and then treated them with intratumoral injections of CAR T cells [71]. In most of their experiments the animals received multiple CAR T-cell injections; however, the authors included data from one mouse that cleared its tumor after receiving only one injection (Fig. 5A). We fit our model to the corresponding longitudinal measurements of tumor size. With appropriate parameter values, our intratumoral injection model accurately simulates the change in tumor size over time, showing tumor eradication around day 15 (Fig. 5B). Without treatment, our model predicts that the tumor would have tripled in size over baseline by day 18. Skovgard et al. assessed the effect of antigen expression density on CAR T-cell kinetics and compared the efficacy of systemic vs. intracavitary delivery of CAR T cells [72]. The authors established antigen-positive orthotopic mesothelioma tumors in mice, and then transferred CAR T cells into the mice via direct pleural injection into the thoracic cavity (Fig. 5D). They monitored CAR T-cell quantity in this group of 5 mice via bioluminescent imaging (BLI) for 8 days following injection. A separate group consisting of 7 mice treated in the same manner was used to measure tumor size via BLI over the same period. We fit our model to the mean value of these groups. Our intracavitary injection model shows tumor eradication by day 4, accompanied by a 6-fold increase in CAR T cells over their baseline measurement, in good agreement with the reported tumor and CAR T-cell levels (Fig. 5E-F). For an explanation of the parameter estimation procedure used to obtain these model fits, see Materials and Methods section 4.5.

**Figure 5:**
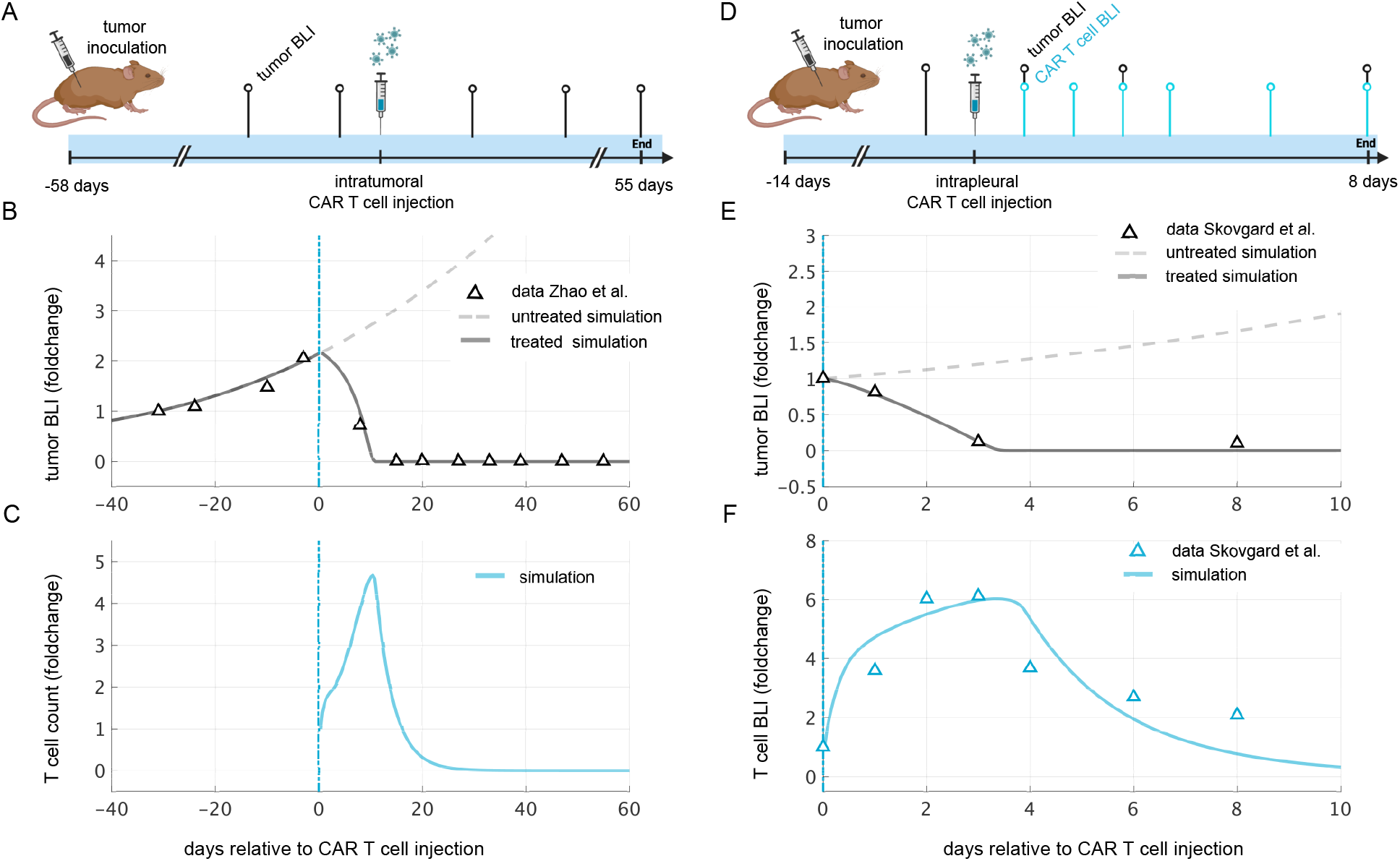
Model fit to data from murine studies. (A) Zhao et al. established large flank mesothelioma tumors in mice, measured baseline tumor size using bioluminescent imaging (BLI), injected 1 × 10^7^ CAR T cells intratumorally 58 days following tumor establishment, and continued to measure the tumor size using BLI for 55 days. (B) Model simulation of untreated tumor and treated tumor overlaid with treated tumor data from Zhao et al. (C) The corresponding CAR T cell count predicted by the model simulation. (D) Skovgard et al. established small antigen-positive orthotopic mesothelioma tumors in mice, measured baseline tumor size using bioluminescent imaging (BLI), injected 1 × 10^7^ CAR T cells intrapleurally 14 days following tumor establishment, and continued to measure the tumor size and CAR T-cell quantity for 8 days using BLI. (E) Model simulation of untreated tumor and treated tumor overlaid with treated tumor BLI quantification from Skovgard et al. (F) The model simulation of CAR T cell count is shown overlaid with CAR T-cell BLI quantification from Skovgard et al. Panels (A) and (D) created with BioRender.com

### 2.6 *in vitro* CAR T-cell exhaustion can result in treatment failure

We next used our model to evaluate the impact of *ex vivo* exhaustion on the efficacy of CAR T-cell therapy given different tumor characteristics and treatment delivery modes. We tested simulated CAR T cells administered either at the center of the tumor or the surface against the four tumor types described in section 2.3 with CAR T-cell doses from 1 × 10^7^ to 1 × 10^9^ cells, spanning a range of reasonable dose levels [27]. Within that dose, we varied the fraction of the CAR T-cell product that is exhausted prior to administration from 0 - 100%, based on analysis by Long et al. [73]. We assumed a detectable diameter of 2 cm at the time of treatment, and simulations were run out to 8 weeks post treatment, at which point the patient response to treatment was classified according to the Response Evaluation Criteria in Solid Tumours (RECIST) [68].

Against all four tumors tested here, intratumoral injection of CAR T cells can eradicate the tumor at high enough doses (Fig. 6 row 1), whereas the best possible outcome from intracavitary delivery is a partial response for tumor types I and II, or stable disease for tumor types III and IV (Fig. 6 row 2). However, otherwise effective doses may fail if the percentage of CAR T cells that are exhausted ex vivo, prior to injection, is too high. For example, an intratumoral injection of 3 × 10^8^ CAR T cells generates a complete response against tumor type IV if 90% of the cells are fully functional, but this same dose is predicted to result in progressive disease if more than 20% of the cells are exhausted (Fig. 6 top right).

**Figure 6:**
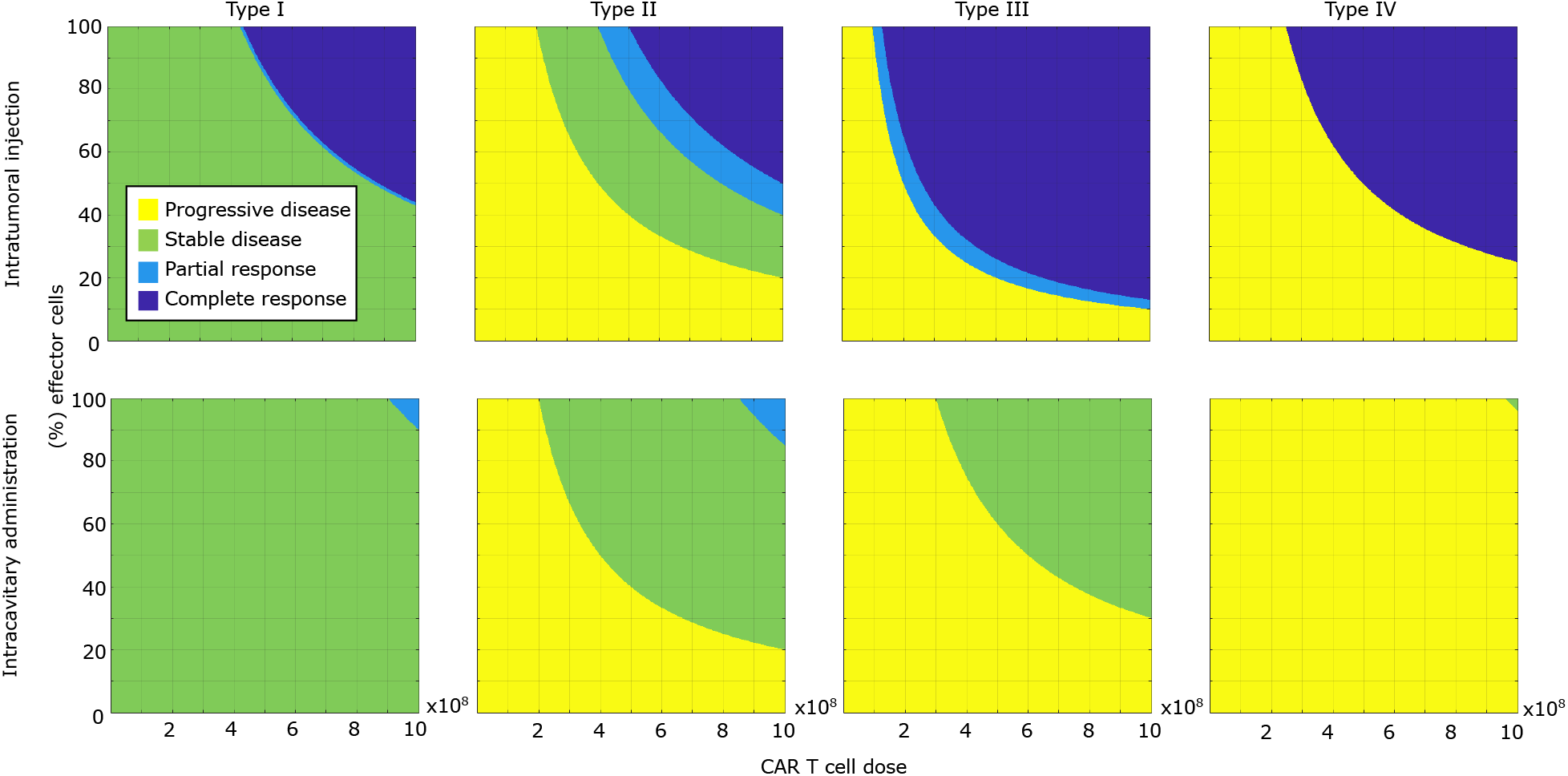
Simulated treatment outcomes vary across different modes of delivery, tumor types, dose levels, and proportions of exhausted cells. Each row corresponds to a mode of local delivery and each column corresponds to a different tumor type as defined in section 4.5, with type I having the longest volume doubling time and type IV having the shortest volume doubling time. Each panel maps a range CAR T cells doses and percentage of non-exhausted cells within that dose to patient outcome, classified using the RECIST criteria[68] at 8 weeks post-treatment. In each simulation, CAR T cell treatment occurred when the detectable tumor radius reached 2 cm.

We also tested the effect of tumor size at the time of CAR T-cell administration on patient outcome by performing these same tests on tumors with detectable diameters of 1 cm and 3 cm at treatment. Our simulations show smaller tumors can be eradicated faster, and with lower doses than larger tumors of the same type (Fig. S2). When treating small tumors, intracavitary administration can induce a complete response for tumor types I-III (Fig. S2). When treating larger tumors, intratumoral injection no longer eradicates tumor type IV even at the highest doses tested (Fig. S2). Note that, unsurprisingly, the timing of evaluation impacts which dose levels are identified as inducing stable disease. Early evaluation can result in a classification of stable disease at lower doses for some scenarios (Fig. S3) and conversely later evaluation requires a higher dose to stabilize tumors, if it is even possible (Fig. S3).

### 2.7 Choice of evaluation criteria impacts classification of partial response and stable disease outcomes

The RECIST criteria for classifying solid tumor response is based solely on the number of tumor lesions and their detectable diameter. Consequently, a treatment that drastically reduces density of the tumor interior but does not impact the extent of the tumor may be falsely classified as completely ineffective (Fig. S4A). As an alternative, we evaluated treatment outcomes for the same scenarios depicted in Fig. 6 using the Choi criteria [74], which assigns a label of partial response if either a density-based or a diameter-based threshold for tumor burden reduction is met (Fig. S4B-C). The minimum dose level to achieve a partial response upon intratumoral injection is lower for all four tumor types, especially the less diffusive tumors, Type I and II (Fig. S4D). For intracavitary administration, the alternate evaluation criteria does not make a significant difference because partial tumor burden reduction occurs at the growth front of the tumor and causes a reduction in detectable tumor diameter.

### 2.8 CAR T-cell treatment fails if CAR T cells move too slowly or too quickly

We performed local sensitivity analysis by varying the value of model parameters one at a time across at least one order of magnitude and simulating treatment of tumor type II with intratumoral injection of 5e8 effector CAR T cells (Fig. S5). For each parameter, we tested a range of values that encompassed successful and unsuccessful treatment. For each simulation, we calculated the minimum tumor cell count following treatment (the tumor nadir), the timing of the tumor nadir, and the maximum number of CAR T cells. We find that there is a non-monotonic relationship between tumor nadir and the CAR T-cell diffusion constant, *D*_*C*_ (Fig. S5A). If CAR T cells diffuse too slowly, CAR T cells kill the core of the tumor but they cannot reach the growth front in order to contain the tumor, and it escapes treatment. However, if the CAR T cells diffuse too rapidly, a substantial quantity leak out of the tumor without engaging and killing tumor cells, leaving the tumor intact. At moderately high diffusion constants, the CAR T cells are able to rapidly spread across the whole tumor but maintain sufficiently high intratumoral concentrations to kill tumor cells and proliferate. Tumor nadir is a monotonic function of the other CAR T parameter values, in agreement with sensitivity analysis of the ODE model studied by Owens and Bozic [45] (Fig. S5B-F). At low values of the maximal CAR T cell killing rate, d, CAR T cells decline without proliferating due to the structure of the model in which proliferation depends on feedback from tumor cell lysis. Initially, as the CAR T-cell killing rate *d* increases, the peak CAR T-cell count is higher and occurs earlier. Further increases in CAR T-cell killing capacity still cause the peak to occur earlier, but actually lower the peak because the tumor is eliminated before high numbers of CAR T cells are generated (Fig. S5B). The CAR T-cell peak was highly sensitive to the maximal CAR T-cell recruitment rate, *j* (Fig. S5E).

## 3 Discussion

In this work we developed a reaction-diffusion model for CAR T-cell treatment of solid tumors and characterized the general behavior of the model in a biologically relevant parameter regime. The underlying mechanism that we use for tumor growth is flexible enough to model a wide range of tumor types. This is particularly valuable as there is significant interest in using CAR T technology to treat diverse cancers. We further demonstrated that this model captures behaviors observed in preclinical trial data by comparing model predictions with data from mouse-imaging studies tracking tumor and CAR T-cell quantities following local delivery of CAR T cells.

We used our reaction-diffusion framework to compare the response of different tumor types to localized CAR T-cell treatment. In particular we considered dense solid tumors with high proliferation but low diffusivity, moderately aggressive solid tumors with both moderate proliferation and moderate diffusivity, highly-diffuse tumors marked by low proliferation but high diffusivity, and aggressive diffusive tumors with both high proliferation and high diffusivity. Surprisingly, the lowest minimum dose was necessary for the low-proliferation, high diffusivity tumors. That is, CAR T-cell therapy was the most effective in treating the tumor with the smallest average density, not the smallest total tumor burden, or the longest VDT. These findings affirm that CAR T cells will not perform equally across different types of solid tumors. It may be particularly promising to pursue CAR T-cell therapy for diffusive tumors. Our analysis suggests that tumor density may be measured when considering the feasibility of CAR T-cell therapy for a particular patient and/or when planning dosages.

There is still a paucity of available data documenting CAR T cell kinetics in human patients following locoregional delivery to treat solid tumors. However, the dynamics of the total number of CAR T cells predicted by our model reflect qualitative features of CAR T cell kinetics observed in murine studies of local delivery, namely an early expansion phase followed by contraction [64, 72]. In our simulations, almost all successful treatments initiated with a reasonable number of CAR T cells display expansion and contraction phases. Only large doses consisting of fully functional cells administered against small tumors are successful without CAR T cell count expanding. Liu et al. reviewed the CAR T cell clinical trial landscape in 2021, which included systemically administered treatments for solid tumors[32]. When comparing responders vs. non-responders across trials, they noted that successful treatments are marked by a higher peak in CAR T cells and a relatively slower contraction rate compared to unsuccessful treatments[32]. Our model simulations align with this observation from clinical data.

We modeled two different modes of local delivery: intratumoral injection vs. intracavitary injection. These options are not always accessible to a patient depending on tumor location and patient health [27]. However, in situations where both are possible, our simulations suggest that intratumoral administration may be successful at lower doses than intracavitary injection. Our results also suggest that for larger tumors, a single dose of CAR T cells administered intracavitarily may induce a partial response or stable disease, but repeat doses of CAR T cells should be explored to move towards tumor eradication. A natural extension of the model would be to optimize the timing of multi-dose regimens.

Using a spatial model is important when studying treatment of solid tumors because assumptions of well-mixed cell populations inherent to previous ODE models do not hold. Furthermore, the movement of CAR T cells within solid tumors has been identified as an important factor in successful treatment. Indeed, Prybutok et al. simulated CAR T-cell therapy using an agent based model and demonstrated that the spatial distribution of tumor cells and healthy cells at the onset of treatment impacted the performance of CAR T cells [75]. Our spatio-temporal model exhibits a failure mode in which there is an initial tumor response to treatment followed by relapse. This clinically observed mode of CAR T-cell failure was not captured by the ODE model from Owens and Bozic [45]. Additionally, as discussed in the above paragraphs, we hypothesize that there may be a potential relationship between tumor density and minimum successful CAR T-cell dose that cannot be captured by an ODE model tracking the total cell populations over time.

This preliminary spatial model has several limitations that could be addressed in future work. A next step with this model framework could be to include the necrotic core observed in larger tumors and compare the outcome of intratumoral injection at different depths. We also assume symmetric spherical growth, but angiogenesis (the development of new blood vessels) is needed to support the growth of a tumor beyond the size of about a million cells, at which point the tumor will no longer develop symmetrically. Because of the lack of conclusive data, we assumed CAR T cells movement is a purely diffusive process. As more data becomes available tracking the distribution of CAR T cells around and within tumors, incorporating chemotaxis or haptotaxis may improve the model. This model considers CAR T cells in effector or exhausted states. However, additional T-cell phenotypes such as regulatory T cells could be explicitly modeled to explore their role in treatment outcomes. This level of granularity would be particularly valuable once there is more data tracking T-cell subsets available for model comparison. Partitioning the CAR T cells into shortlived effector cells and long-lived memory cells may also improve the ability to capture long-term persistence. In clinical data, CAR T cells tend to persist at about ∼ 10% of their peak level [32, 76], but in our model they drop to non-detectable levels. However, our model does reflect the fact that CAR T-cell persistence is important to successful treatment. Increasing the CAR T-cell death or inactivation rate causes treatment to lose effectiveness, and conversely lowering these parameters causes treatment to be more effective.

CAR T cell therapy for solid tumors faces numerous challenges, but significant progress is being made. This work lays out an accessible mathematical modeling framework for studying one possible advance: locoregional delivery of CAR T-cells. Simulation results suggest several testable hypothesis and the model is ready for extension and refinement as more data on CAR T-cell dynamics becomes available.

## 4 Materials and Methods

### 4.2 Mathematical modeling

We use a reaction-diffusion model to describe changes in cell density over time. The density of tumor cells and CAR T cells at location *x* and time *t* are denoted by *u*(*x, t*) and *v*(*x, t*) respectively. The rate of change in the density of the tumor is the sum of density-dependent diffusion, and a forcing term *F*_1_ that accounts for tumor growth in the absence of treatment and for the effect of CAR T cells on the tumor [45]. We write the tumor concentration as

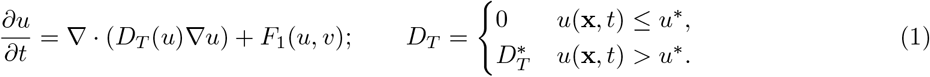

Thus diffusion does not occur below a given critical tumor concentration, *u*^*^, but above the critical concentration cells diffuse at a constant rate,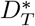. A Dirichlet boundary condition of u = 0 at infinity is enforced, allowing unrestricted tumor expansion. Further details on boundary conditions of the PDE are included in the supplementary material. The evolution of the CAR T cells is a diffusive process with the addition of a forcing function, *F*_2_, that describes the interactions between tumor cells and CAR T cells [45]:

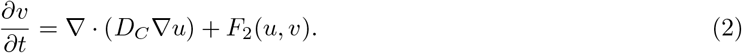

A Neumann boundary condition is used to allow for proliferation at the edge of the tumor region without precipitous leakage, the details of which are reported in the supplementary material. We constructed the forcing functions, *F*_1_ and *F*_2_, by adapting the ODE model studied by Owens and Bozic [45]. Let

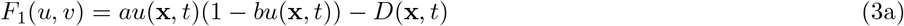

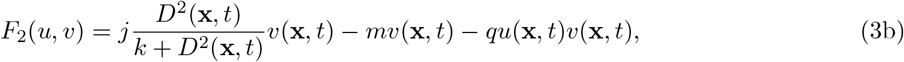

Where

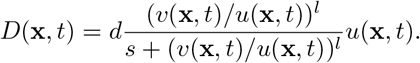

The proliferation term for tumor cells in our model was drawn directly from the ODE model for CAR T-cell therapy from Owens and Bozic [45]. In the absence of CAR T cells, the proliferation of tumor cells is driven by logistic growth. Previous reaction-diffusion models of tumor growth have considered exponential, gompertzian, and logistic growth [53]. Tumor cells are killed by CAR T cells in a saturating density-dependent manner that depends on the ratio of tumor to CAR T-cell density in a given location. The proliferation terms governing CAR T cells are adapted slightly from the Owens and Bozic model [45]. Modifications to the recruitment term were required because we do not include endogenous effector cells in this model. In the resulting forcing function, CAR T-cell recruitment is driven by cell lysis, while CAR T-cell density decreases due to cell death and exhaustion upon repeated interaction with tumor cells.

In this work, we consider the evolution of a radially symmetric, spherical tumor growing on an infinite domain. This approach was also taken by Burgess et al. in their 3 dimensional extension of the work of Swanson et al. modelling glioma [54]. Applying this geometric consideration to equations (1) and (2) and keeping the boundary conditions as is, yields

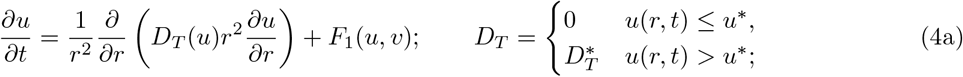

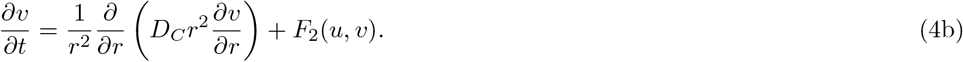

The nondimensional version of the model is derived in the supplementary material. Our model incorporates the conversion of effector CAR T cells to an exhausted state through a mass-action exhaustion term in the CAR T-cell forcing function, −*quv*. In our reaction-diffusion model, CAR T cells are assumed to have an effector phenotype, hence all CAR T cells contribute to antitumor activity. However, experimental measurements of CAR T-cell kinetics over time do not discriminate between phenotypes. In order to compare simulation results with data, we add an ODE to the model to track the exhausted CAR T cells as well. Assuming that the exhausted CAR T cells have the same death rate as the effector CAR T cells, *m*, and an initial number of cells *v*_*E*_(0), the number of exhausted CAR T cells at time *t* is given by

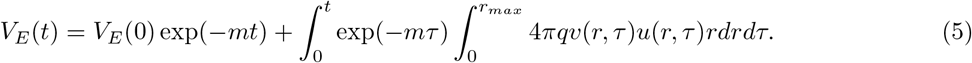

The total number of CAR T cells at time *t* can then be calculated by summing the exhausted and the effector cells.

### 4.2 Parameter Estimation

In order to carry out informative numerical analysis and simulations of the model, we estimated several parameter values from experimental data and identified others from previous modeling work. We consider diffusion constants for tumor density in a range of 10^−5^ − 10^−4^ cm^2^/day. The upper end of this range is used to model diffusive tumors, and is calculated from measurements of the velocity of individual glioma cells in vitro [77]. The diffusion constants for more compact tumors are an order of magnitude smaller, aligning with the range reported by Weis et al., who calibrated their reaction diffusion equation to breast cancer imaging data [78]. As the local carrying capacity for tumor cells, we use 2.39 × 10^8^ cells/cm^3^, based on assuming an average cell radius of 10 *µ*m, as in previous reaction diffusion models of tumor growth by Swanson et al. [79, 80]. The density above which diffusion of tumor cells occurs ranges from 1-50% of the local carrying capacity, allowing for a wide range of tumor density profiles. We use 10^8^ cells/cm^3^ as the lower limit of detection when determining tumor radius for generating initial tumor conditions and evaluating the efficacy of treatment. This detection threshold is around 40% of the carrying capacity, which falls between the values used by Swanson et al. to model a highly sensitive imaging modality (16% of carrying capacity) and a less sensitive modality (80% of carrying capacity) [56]. The tumor proliferation rates studied here range from 0.025 − 0.25 day^−1^, on the same order as previous models of tumor growth [45, 54, 55, 78]. Particular values were selected to achieve tumor volume doubling times relevant to malignant gliomas (15-21 days) [67], breast cancer (46-825 days) [66], and liver cancer (85-150 days) [65] with a focus on the more aggressive end of each range because slow growing tumors are unlikely candidates for CAR T-cell therapy. We computed the diffusion constant for CAR T cells, *D*_*C*_ = 1.38 cm^2^/day based on intratumoral CAR T-cell velocities reported from murine imaging studies by Mullazzani et al. [64]. The parameters dictating CAR T-cell proliferation, exhaustion, clearance, and killing of tumor cells came from the Patient 4 parameter set, reported in Owens and Bozic [45]. To reflect the reduced efficacy of CAR T cells against solid tumors compared with blood cancers, we scaled these parameters by 20%. See tables S1 and S2 for a complete list of parameter sets used in model simulations.

### 4.3 Numerical analysis

To integrate the PDE given by (4) we employ Crank-Nicolson [81] for both the tumor and CAR T cell densities. The standard Crank-Nicolson scheme for the homogeneous diffusion equation,

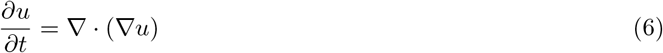

has an order of convergence of *o* (Δ*t*^2^, Δ*r*^2^); i.e., second order in time and space. However, the PDE in the present study, (4), would not be expected to maintain the same order of accuracy due to the sharp boundary between proliferative and diffusive regions. Using the method of manufactured solutions we observe a local order of accuracy at *r* = 0.5 of *o* (Δ*t*^1.5^, Δ*r*^1.1^) and *o* (Δ*t*^1^, Δ*r*^0.6^) for *u* and *v* respectively. Towards the boundary of the computational domain the error increases and we observe a maximum order of accuracy of *o* (Δ*t*^0.9^, Δ*r*^1^) and *o*(Δ*t*^0.3^, Δ*r*^0.5^) for *u* and *v* respectively. Since the agorithm performs much better towards the center of the computational domain, we choose a domain large enough to keep the extent of the tumor well within the boundary of the domain. The CAR T cells, unlike tumor cells, are free to spread out, but in the absence of tumor cells they no longer proliferate. Consequently, the density of CAR T cells far from the tumor is low in model simulations, as observed in murine experiments. Therefore, although we may detect CAR T-cells towards the boundary of the computational domain, their numbers are several orders of magnitude less than around the tumor, and even the CAR T-cells near the computational boundary experience an error less than 10% compared to the ground truth according to the manufactured solution. For further details on the numerical analysis see the supplementary material.

### 4.4 Numerical simulations

All results were generated from simulations implemented with MATLAB R2021b. We used a spatial grid with step size Δ*r* = 0.015 cm and time step Δ*t* = min(*a*Δ*r/D*_*C*_, Δ*r/*5) days. This condition for the time step was empirically chosen to balance efficiency and numerical stability. The extent of the spatial domain was selected based on the initial tumor size and anticipated tumor growth such that the tumor would not reach within 1 cm of the edge of the domain. All model parameter values used in simulations are included in supplementary tables S1 and S2.

### 4.5 Process for fitting model to murine data

We digitized data from two studies on locoregional delivery of CAR T cells using WebPlotDigitizer [82]. The first set of data comes from Figure 4c in Zhao et al. [71], which reports tumor burden prior to and after treatment with a single intratumoral dose of CAR T cells. We fixed the carrying capacity and diffusion threshold for tumor cells, then estimated the tumor growth rate and diffusion constant to minimize the least squares error when comparing the total tumor burden predicted by the model to the pre-treatment data points. We fixed the CAR T-cell diffusion constant at *D*_*C*_ = 0.0138 cm^2^/day as estimated based on Mullazanni et al. We fixed all remaining CAR T-cell parameters except the maximal lysis rate, *d*, at the values derived from Patient 3 in Owens et al. [45]. We started with an initial guess of *d* = 2.25, from Patient 3 in Owens et al., and increased the value of *d* until the tumor burden was reduced sufficiently quickly to match the data.

The second set of data came from Figure 4F in Skovgard et al., which reports tumor and CAR T cell levels measured via BLI following treatment with a single intrapleural dose of CAR T cells. [72]. We specifically extracted the mean values for each cell population. In this case, because there are no pre-treatment tumor measurements, we fixed the tumor parameters around those used to define tumor type IV. We fixed the CAR T-cell diffusion constant, the lysis rate for half maximal CAR T-cell proliferation, the CAR T-cell exhaustion rate, and the CAR T-cell death rate at the same values used for the simulation of Zhao et al. data. For the remaining 4 CAR T-cell parameters, we performed an iterative search to identify a set of values that fit both the tumor and the CAR T-cell data well. We started with initial parameter ranges reported in table S2, then ran simulations with values randomly selected from the parameter space using latin hypercube sampling. We re-centered and narrowed the parameter ranges around the average value from the best 3 runs, and repeated this process until a satisfactory fit to the data was achieved.

## Data and code availability

Murine data used in this paper was previously published by Skovgard et al. [72] and Zhao et al.[71]. The digitized values used for model parameter estimation are available in the supporting data file. MATLAB code used to produce the results included in this work is available on github at https://github.com/lacyk3/RadiallySymmetricCART.

## Author contributions and disclosures

K.O., A.R., and I.B. designed and performed research; A.R. and K.O. wrote software; and K.O., A.R., and I.B. wrote the paper. The authors have no competing interests to disclose.

## Acknowledgements

This work is supported by the WiSTEM^2^D Scholar Award from Johnson & Johnson and by the National Institute of Allergy and Infectious Diseases of the NIH award T32AI118690. The content is solely the responsibility of the authors and does not necessarily represent the official views of the NIH.

