## Supplementary Information for "Spatiotemporal dynamics of tumor - CAR T-cell interaction following local administration in solid cancers"

This PDF file includes:

- Supporting text
- Figures S1 to S6
- Tables S1 to S2
- SI References

### S1 Nondimensionalization

We nondimensionalize Eq. (4) with $\tau = at$, $\hat{r} = r/R_0$, $\hat{u} = bu$, and $\hat{v} = bv$, where $a$ is the tumor proliferation rate from [1], $R_0$ is the initial spread of the tumor, and $1/b$ is the maximum tumor cell density. This gives us the equation (with hats removed)

$$\frac{\partial u}{\partial t} = \frac{1}{r^2}\frac{\partial}{\partial r}\left(D_T(u) r^2 \frac{\partial u}{\partial r}\right) + F_1(u,v); \qquad D(u) = \begin{cases} 0 & u(r,t) \le \hat{u}^*, \\ D_T^* & u(r,t) > \hat{u}^*; \end{cases} \tag{S1a}$$

$$\frac{\partial v}{\partial t} = \frac{1}{r^2}\frac{\partial}{\partial r}\left(D_C r^2 \frac{\partial v}{\partial r}\right) + F_2(u,v). \tag{S1b}$$

with

$$F_1 = \left[1 - u - \gamma \frac{v^l}{su^l + v^l}\right] u(r,t) \tag{S2a}$$

$$F_2 = \alpha \left[\frac{v^{2l}u^2}{(su^l + v^l)^2 + \zeta_C v^{2l} u^2} - \beta_C u - \chi\right] v(r,t) \tag{S2b}$$

where $\hat{u}^* = bu^*$, $\alpha = d^2 j/kab^2$, $\beta_C = kqb/d^2 j$, $\gamma = d/a$, $\chi = mkb^2/d^2 j$, $\zeta = d^2/kb^2$, $s = 0.25$, and $l = 1.36$.

### S2 Numerical Methods

Finite difference schemes are employed to solve the radially symmetric spherical diffusion equations in the model (4). Namely, Crank-Nicolson (CN) [2] is used for the results presented in the body of the paper. For the concentration of CAR T cells, $v_m^n = v(m\Delta r, n\Delta t)$, the CN scheme is fairly standard yielding

$$\begin{aligned} v_m^{n+1} - v_m^n - \frac{D_C}{r_m}\left[v_{m+1}^{n+1} - v_{m-1}^{n+1} + v_{m+1}^n - v_{m-1}^n\right]\frac{\Delta t}{2\Delta r} \\ - D_C\left[v_{m+1}^{n+1} - 2v_m^{n+1} + v_{m-1}^{n+1} + v_{m+1}^n - 2v_m^n + v_{m-1}^n\right]\frac{\Delta t}{2(\Delta r)^2} \\ - \left[F_2\left(u_m^{n+1}, v_m^{n+1}\right) + F_2\left(u_m^n, v_m^n\right)\right]\frac{\Delta t}{2} = 0, \end{aligned} \tag{S3}$$

with $v_{m+1} = v_{m-1}$ at $r = 0$ and for $r \in \partial\Omega(u)$

For the tumor concentration, $u_m^n = u(m\Delta r, n\Delta t)$, the diffusivity, $D_T$, depends on $u$, which means its dependence on $r$ is changing in time. If $u_{m+1}^n$ and $u_{m-1}^n$ are both larger than $\hat{u}^*$ or both less than $\hat{u}^*$, the central difference for $D_T$ is trivial, and hence $D_T$ is treated as a constant. However, if the parity is different, there is no flux from the side with a concentration less than $\hat{u}^*$, and therefore it can be treated as a Neumann boundary. If $u_{m+1}^n$ and $u_{m-1}^n$ have the same parity relative to $\hat{u}^*$ we write

$$\begin{aligned} u_m^{n+1} - u_m^n - \frac{D_T}{r_m}\left[u_{m+1}^{n+1} - u_{m-1}^{n+1} + u_{m+1}^n - u_{m-1}^n\right]\frac{\Delta t}{2\Delta r} \\ - D_T\left[u_{m+1}^{n+1} - 2u_m^{n+1} + u_{m-1}^{n+1} + u_{m+1}^n - 2u_m^n + u_{m-1}^n\right]\frac{\Delta t}{2(\Delta r)^2} \\ - \left[F_1\left(u_m^{n+1}, v_m^{n+1}\right) + F_1\left(u_m^n, v_m^n\right)\right]\frac{\Delta t}{2} = 0, \end{aligned} \tag{S4}$$

otherwise we write

$$u_m^{n+1} - u_m^n - D_T\left[u_\pm^{n+1} - u_m^{n+1} + u_\pm^n - u_m^n\right]\frac{\Delta t}{(\Delta r)^2} - \left[F_1\left(u_m^{n+1}, v_m^{n+1}\right) + F_1\left(u_m^n, v_m^n\right)\right]\frac{\Delta t}{2} = 0, \tag{S5}$$

where $u_\pm = u_{m+1}^n$ if $u_{m+1}^n > \hat{u}^*$ or $r_m = 0$, and $u_\pm = u_{m-1}^n$ if $u_{m-1}^n < \hat{u}^*$.

#### S2.1 Boundary and Initial Conditions

In this section consider $\partial\Omega(u) = 4\max(R_{\text{tumor}})$ to be the boundary of the region of influence of the tumor $\Omega$, where $R_{\text{tumor}}$ are the radial values such that $u > 0$; that is, only active CAR T cells within the influence of the tumor region are considered since most CAR T cells remain near the tumor.

Since diffusion is the main driver of the dynamics before cell-to-cell interactions begin, we consider an initial profile with a compact Gaussian shape (i.e., a bump function) shortly after injection. The bump function assumption simplifies the model by obviating the dynamics of the injection itself. Both the internal and external injection bump functions are illustrated in Fig. 1.

At the boundary, $\partial\Omega$, we consider a Neumann condition

$$\frac{\partial u}{\partial r} = \frac{\partial v}{\partial r} = 0, \tag{S6}$$

since we keep our boundary away from the tumor and we want to prevent precipitous leakage of the CAR T-cells because CAR T-cells tend to stay close to the tumor. To implement the condition at the boundary, we use ghost points, where for the purpose of the centered difference, we assume just past the boundary and just before the boundary the concentration is equivalent. That is, if $m = M$ corresponds to $r = \partial\Omega$, then $u_{M+1} = u_{M-1}$ and $v_{M+1} = v_{M-1}$. This yields the equations

$$u_M^{n+1} - u_M^n - D_T\left[u_{M-1}^{n+1} - u_M^{n+1} + u_{M-1}^n - u_M^n\right]\frac{\Delta t}{(\Delta r)^2} - \left[F_1\left(u_M^{n+1}, v_M^{n+1}\right) + F_1\left(u_M^n, v_M^n\right)\right]\frac{\Delta t}{2} = 0, \tag{S7a}$$

$$v_M^{n+1} - v_M^n - D_T\left[v_{M-1}^{n+1} - v_M^{n+1} + v_{M-1}^n - v_M^n\right]\frac{\Delta t}{(\Delta r)^2} - \left[F_1\left(u_M^{n+1}, v_M^{n+1}\right) + F_1\left(u_M^n, v_M^n\right)\right]\frac{\Delta t}{2} = 0. \tag{S7b}$$

#### S2.2 Manufactured Solution

Now let us consider the convergence properties using the manufactured solution

$$\tilde{u}(r,t) = \frac{1}{r}e^{-D_T^* t}\sin(r) \tag{S8a}$$

$$\tilde{v}(r,t) = \frac{1}{r}e^{-D_C t}\sin(r) \tag{S8b}$$

to test $D_T(u) = D_T^*$ This yields the test PDEs

$$\frac{\partial u}{\partial t} = \frac{1}{r^2}\frac{\partial}{\partial r}\left(D_T^* r^2 \frac{\partial u}{\partial r}\right) + F_1(u,v) - F_1(\tilde{u},\tilde{v}); \tag{S9a}$$

$$\frac{\partial v}{\partial t} = \frac{1}{r^2}\frac{\partial}{\partial r}\left(D_C r^2 \frac{\partial v}{\partial r}\right) + F_2(u,v) - F_2(\tilde{u},\tilde{v}) \tag{S9b}$$

We employ the Crank-Nicolson scheme from Sec. 4.3 on the test PDE (S9) to get the approximate solution $u(r,t)$ and $v(r,t)$. Then using the ground truth (S8), we find the absolute errors $|u(r,t) - \tilde{u}(r,t)|$ and $|v(r,t) - \tilde{v}(r,t)|$. In Fig. S7, we illustrate the error in practice and the convergence properties of the method for our setup. As our PDE, and therefore the test PDE as well, has a sharp boundary between the proliferative and diffusive regimes in addition to significant nonlinearities, we would not expect our implementation of the Crank-Nicolson finite difference scheme to have the same convergence properties as that of the standard diffusion equation. Even still, as we observe in Fig. S7, the error is quite low especially near the center of the domain where it is nearly $10^{-4}$. Towards the boundary of the computational domain the error does increase to slightly greater than 1%, however we can mitigate this issue by expanding our computational domain to keep the tumor away from the boundary. As we expect, the convergence properties away from the boundary are closer to that of the standard diffusion equation with local order of convergence of $o\left(\Delta t^{1.5}, \Delta r^{1.1}\right)$ and $o\left(\Delta t^{1}, \Delta r^{0.6}\right)$ for $u$ and $v$ respectively.

#### S3 Supplementary Figures

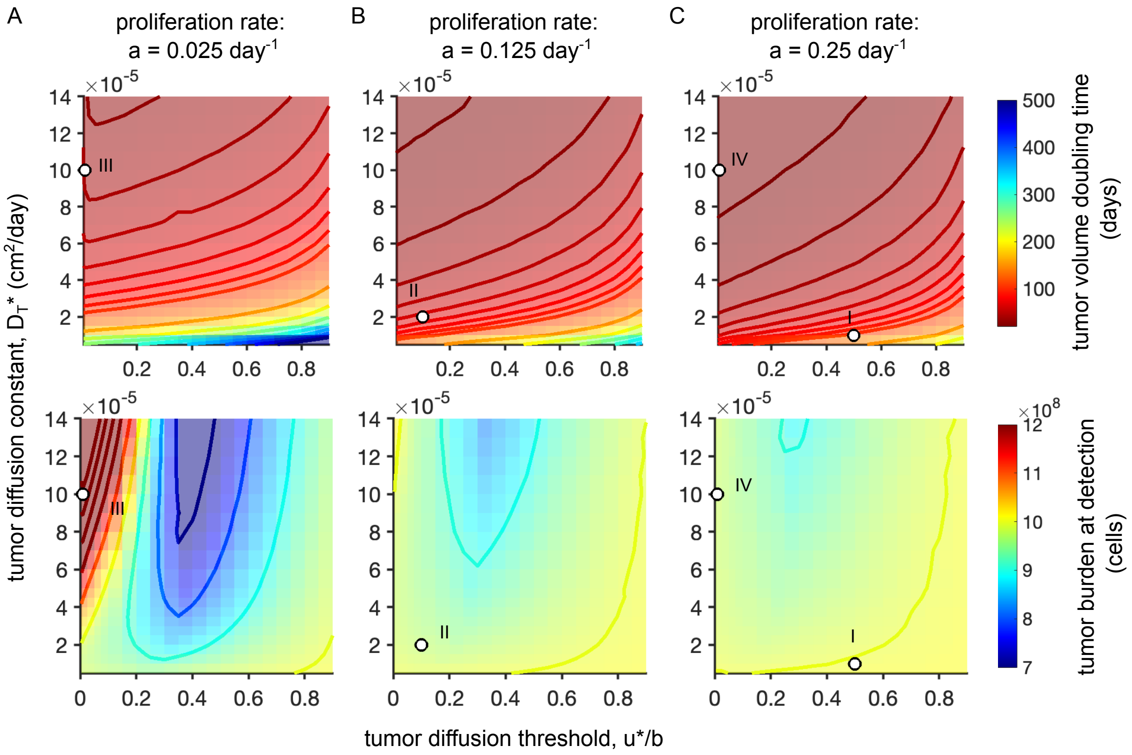


Figure S1: *Model-predicted tumor volume doubling time (VDT) and tumor burden at detection as a function of tumor growth parameters.* In each column a fixed value was used for the tumor cell proliferation rate, while the tumor diffusion constant was varied from $5e-6$ to $1.4e-4$ cm$^2$/day and the threshold tumor density governing the onset of diffusion was varied from 1-90% of the tumor density carrying capacity. Panel A shows tumor behavior at a low proliferation rate of $a = 0.025$ day$^{-1}$, panel B a medium proliferation rate of $a = 0.125$ day$^{-1}$, and panel C a high proliferation rate, $a = 0.25$ day$^{-1}$. For each simulation we calculated VDT (top row) by recording the time a spherical tumor grew from a detectable radius of 1 cm to a detectable radius of $2^{1/3}$ cm. We also calculated the tumor burden at detection by integrating the tumor cell density across the spatial domain to get total cell count when the detectable diameter was 2 cm. The tumor growth parameters defining tumor types I-IV are indicated by white circles on the appropriate panel.

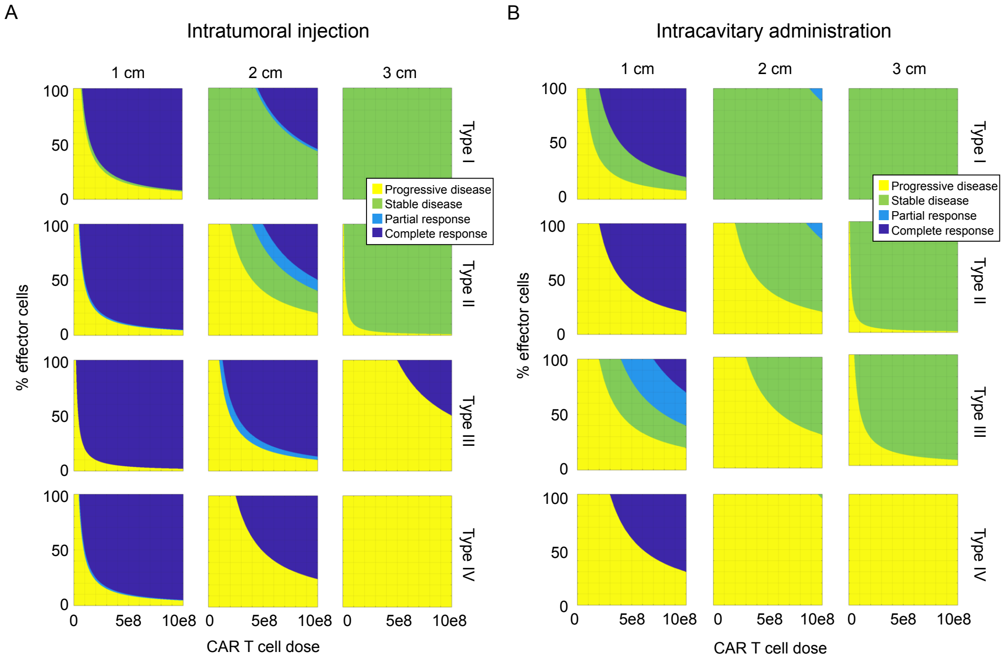


Figure S2: *Outcome maps given varied tumor size at the time of treatment.* We simulated (A) intratumoral injection and (B) intracavitary administration of CAR T cells to treat tumors with varied growth parameters and initial sizes. Within each subfigure, the row corresponds to a different tumor type as defined in section 4.5, with type I having the longest volume doubling time and type IV having the shortest volume doubling time. The columns correspond to a different tumor radius at the time of treatment with column 1 being 1 cm, column 2 is 2 cm and column 3 is 3 cm. Each panel maps a range CAR T cells doses and percentage of non-exhausted cells within that dose to patient outcome, classified using the RECIST criteria[3] at 8 weeks post-treatment.

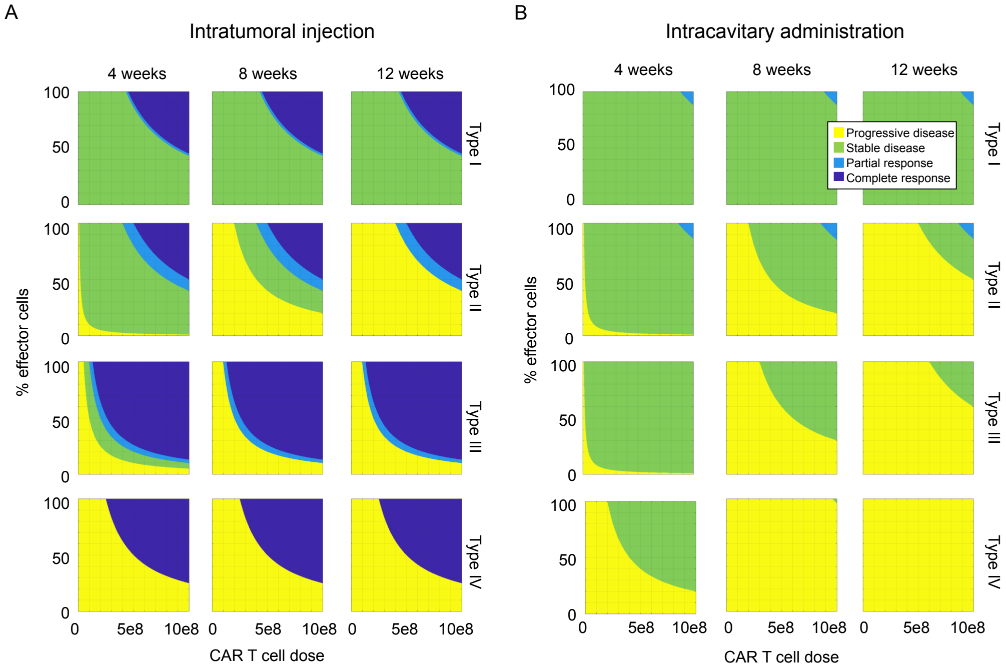


Figure S3: *Outcome maps given varied time of patient evaluation.* We simulated (A) intratumoral injection and (B) intracavitary administration of CAR T cells to treat varied tumor types. Within each subfigure, the row corresponds to a different tumor type as defined in section 2.2, with type I having the longest volume doubling time and type IV having the shortest volume doubling time. The columns within each subfigure correspond to a different date of evaluation, with column 1 being evaluation at 4 weeks, column 2 at 8 weeks, and column 3 at 12 weeks post-treatment. Each panel maps a range CAR T cells doses and percentage of non-exhausted cells within that dose to patient outcome. In each simulation, CAR T cell treatment occurred when the detectable tumor radius reached 2 cm.

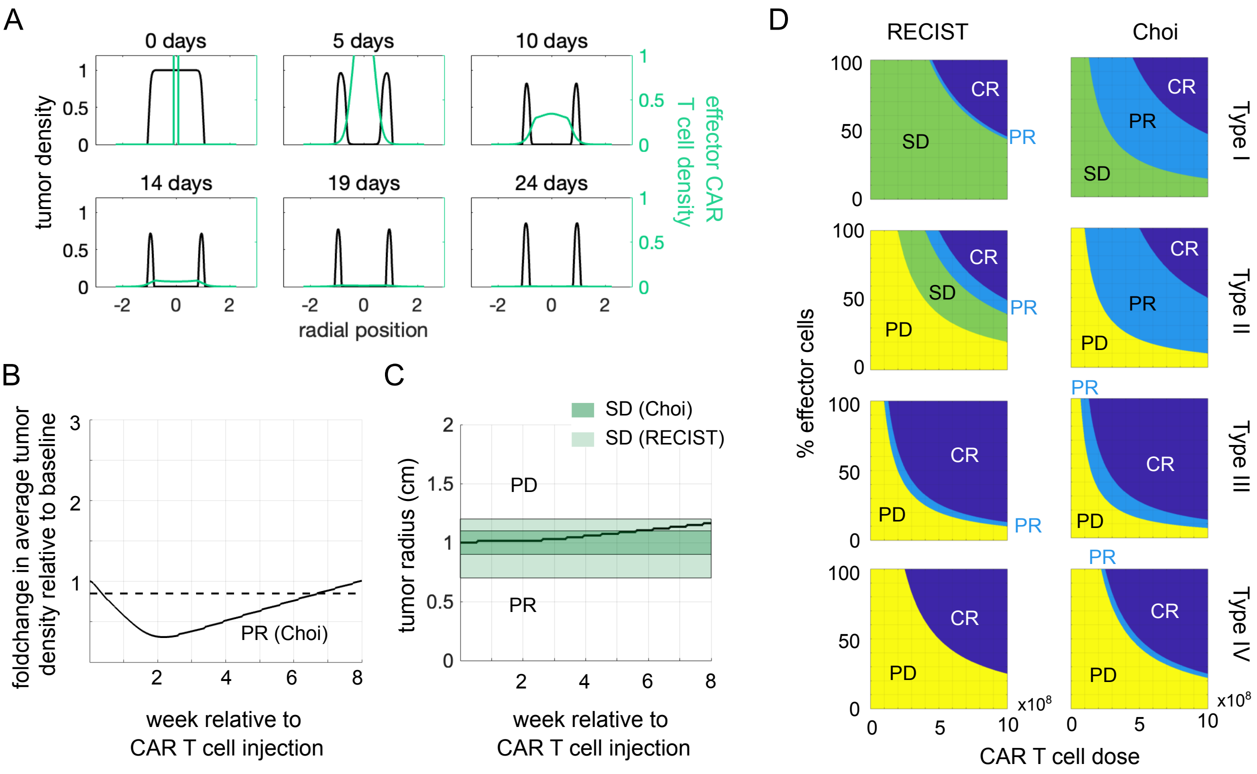


Figure S4: *Classification of treatment outcome following intratumoral CAR T cell treatment evaluated using Choi versus RECIST criteria.* (A) Tumor and effector CAR T cell densities over the first 24 days following intratumoral injection of $3 \times 10^8$ effector CAR T cells to treat a 2 cm diameter tumor of type II show wide variation across the spatial domain. (B) The foldchange in average tumor density over time reflects the effect of CAR T cells killing tumor cells at the tumor core. Because tumor burden drops below the horizontal dotted line at 0.85, the outcome is classified as partial response to therapy by the Choi criteria. (C) The detectable tumor burden for the same scenario increased monotonically. The size-based criteria for stable disease according the Choi criteria requires staying in the region highlighted in dark green, which this scenario does not meet. The boundaries for stable disease according to the RECIST criteria are highlighted in light green, which this scenario does meet. (D) Outcome maps if four tumor types (row 1-4) were treated with intratumoral injeciton of a range CAR T cells doses and percentage of non-exhausted cells within that dose, and evaluated at 8 weeks post-injection via the RECIST criteria (column 1) or the Choi criteria (column 2).

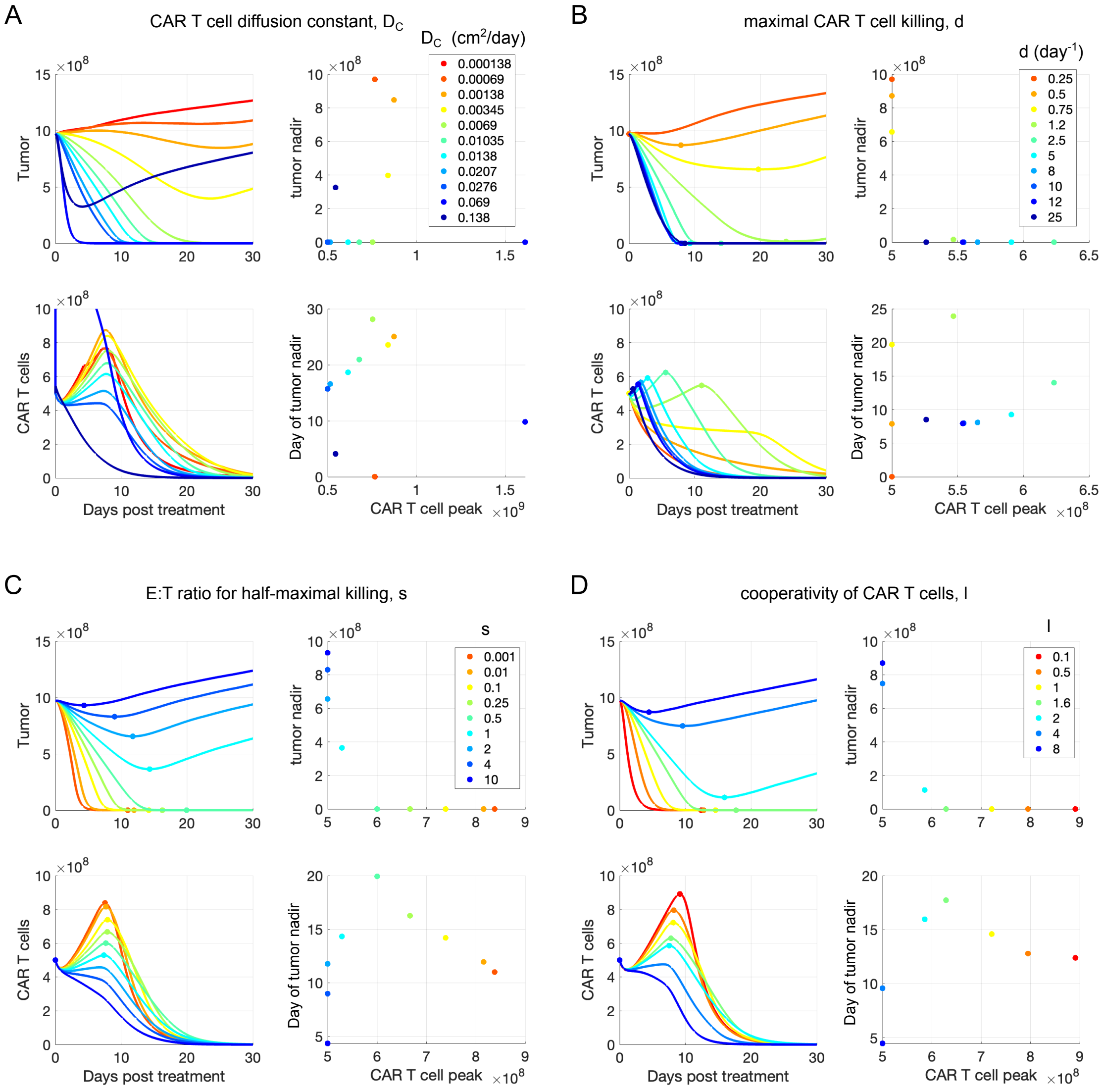


Figure S5: *Tumor and CAR T cell trajectories for varied values of CAR T cell parameters.* We varied each model parameter associated with CAR T cell behavior one at a time across a range of values and simulated intratumoral injection of $5 \times 10^8$ CAR T cells to treat tumor type II out to 30 days post-injection. The resulting trajectories for total tumor cell count and total CAR T cell count are shown, colored according to the value of the parameter being studied. For each trajectory, we plot the minimum tumor cell count following injection, called the tumor nadir, and the time of tumor nadir as functions of the peak number of CAR T cells. This figure illustrates the impact of (A) the CAR T cell diffusion constant, $D_C$, (B) the maximal CAR T cell killing rate, $d$, (C) the ratio of T cells to tumor cells for half-maximal killing, $s$, and (D) the cooperativity of CAR T cell killing, $l$. See Figure S6 for remaining parameters.

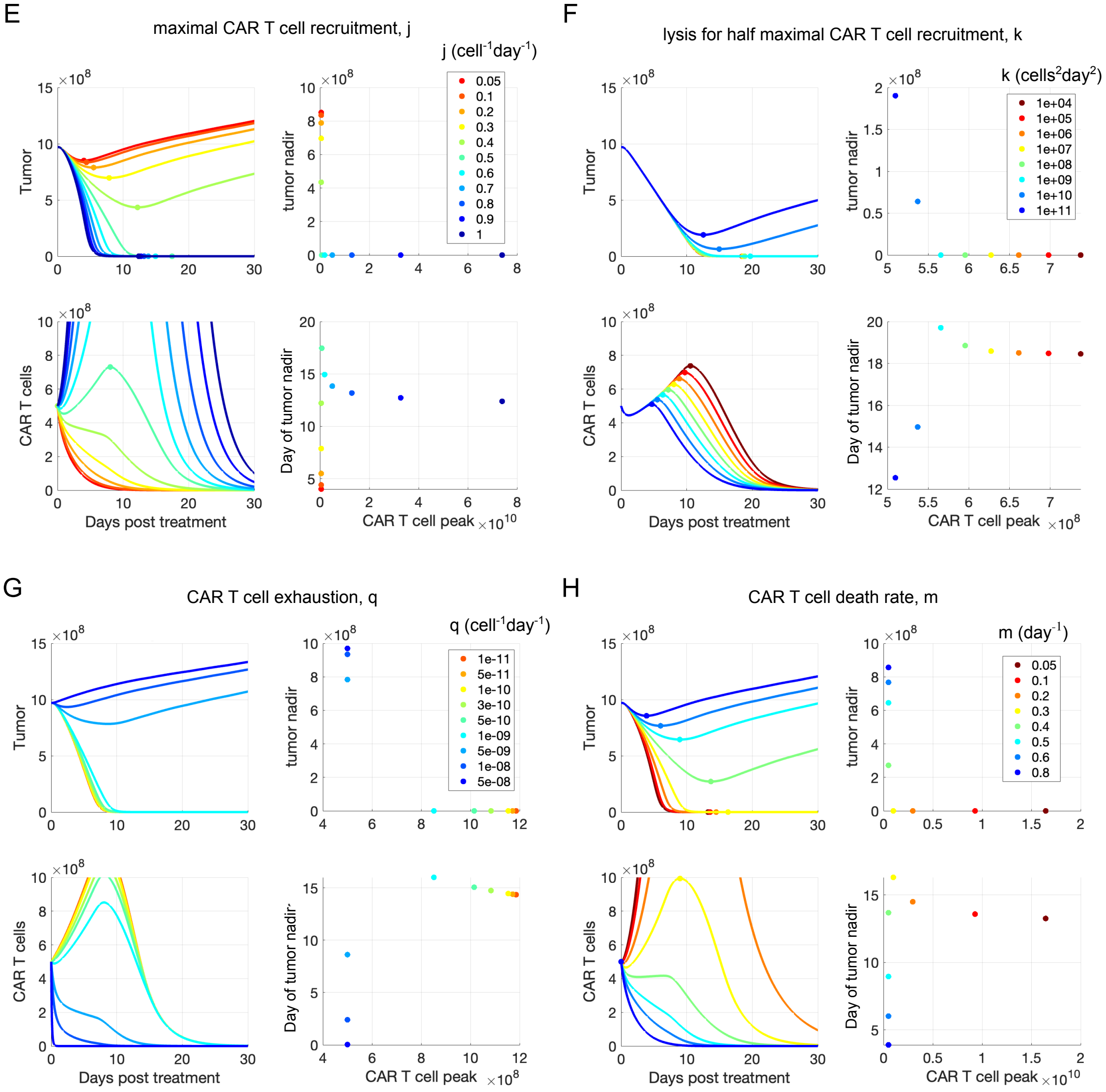


Figure S6: *Tumor and CAR T cell trajectories for varied values of CAR T cell parameters (continuted).* (E) the maximal CAR T cell proliferation/recruitment rate, $j$, (F) the lysis rate for half maximal CAR T cell expansion, $k$, (G) the CAR T cell exhaustion rate, $q$, and (D) the CAR T cell death rate, $m$.

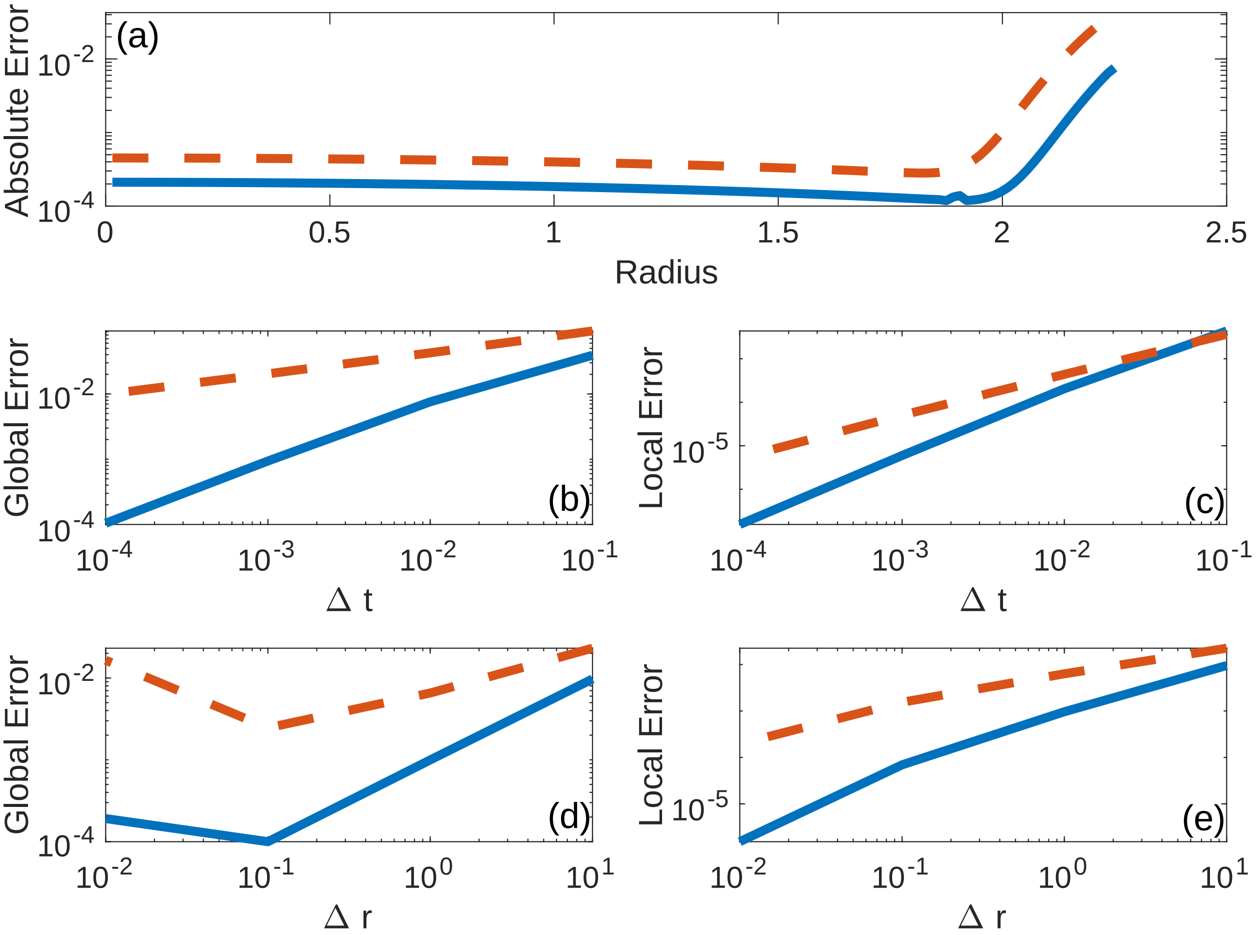


Figure S7: *Convergence analysis via method of manufactured solutions.* Dashed (red) curve indicates $v(r,t)$ and solid (blue) curves indicate $u(r,t)$. **(a)** Absolute spatial error for $\Delta t = 0.01$ and $\Delta r = 0.015$ for $t \gg 0$. **(b)** Absolute global error for $\Delta t = 0.1, 0.01, 0.001, 0.0001$. **(c)** Absolute local error for $\Delta t = 0.1, 0.01, 0.001, 0.0001$ at $r = 0.5$. **(d)** Absolute global error for $\Delta r = 10, 1, 0.1, 0.01$. **(e)** Absolute local error for $\Delta r = 10, 1, 0.1, 0.01$ at $r = 0.5$.

#### S4 Supplementary Tables

| symbol | description | unit | Type I | Type II | Type III | Type IV |
|---|---|---|---|---|---|---|
| $a$ | tumor proliferation rate | $\text{day}^{-1}$ | 0.25 [4, 5] | 0.125[4] | 0.025[6] | 0.25 [6] |
| $D_{T^*}$ | tumor diffusion constant | $\text{cm}^2/\text{day}$ | 1e-5 [7] | 2e-5 [7] | 1e-4 [8] | 1e-4 [8] |
| $u^*$ | tumor diffusion threshold | $\text{cells/cm}^3$ | 1.2e8 | 2.39e7 | 2.39e6 | 2.39e6 |
| $1/b$ | tumor carrying capacity | $\text{cells/cm}^3$ | 2.39e8 [9] | | | |
| $D_C$ | CAR T cell diffusion constant | $\text{cm}^2/\text{day}$ | 1.38e-2 [10] | | | |
| $d$ | maximum lysis rate | $\text{day}^{-1}$ | 1.8 [1] | | | |
| $s$ | E:T ratio for half-maximal lysis | unitless | 0.55 [1] | | | |
| $l$ | CAR T cell cooperativity | unitless | 1.7 [1] | | | |
| $j$ | maximum CAR T cell proliferation rate | $\text{day}^{-1}$ | 0.48 [1] | | | |
| $k$ | lysis rate for half-maximal CAR T cell proliferation | $\text{cells}^2/\text{day}^2$ | 2.4e7 [1] | | | |
| $q$ | CAR T cell exhaustion rate | $\text{cells}^{-1}\text{day}^{-1}$ | 1.84e-9 [1] | | | |
| $m$ | CAR T cell death rate | $\text{day}^{-1}$ | 0.35 [1] | | | |

Table S1: Parameter values used in numerical simulations of the 4 tumor types described in section 2.2. The source used to determine a reasonable value for each parameter is indicated in brackets next to each value. A fraction of the tumor carrying capacity was chosen as the tumor diffusion threshold for each tumor type to obtain characteristic, qualitative behavior. For parameter values below the horizontal line (tumor carrying capacity and CAR T cell related parameters), the same values were used when simulating all tumor types.

| symbol | description | unit | Zhao et al. | Skovgard et al. | range tested |
|---|---|---|---|---|---|
| $a$ | tumor proliferation rate | day$^{-1}$ | 0.08 | 0.2 | |
| $D_{T^*}$ | tumor diffusion constant | cm$^2$/day | 8e-5 | 1e-4 | |
| $u^*$ | tumor diffusion threshold | cells/cm$^3$ | 2.39e-7 | 2.39e-7 | |
| $1/b$ | tumor carrying capacity | cells/cm$^3$ | 2.39e8 | 2.39e8 | |
| $D_C$ | CAR T cell diffusion constant | cm$^2$/day | 1.38e-2 | 1.38e-2 | |
| $d$ | maximum lysis rate | day$^{-1}$ | 12 | 20 | 0.1-25 |
| $s$ | E:T ratio for half-maximal lysis | unitless | 0.25 | 0.2 | 0.1-10 |
| $l$ | CAR T cell cooperativity | unitless | 1.36 | 1.24 | 0.1-4 |
| $j$ | maximum CAR T cell proliferation rate | day$^{-1}$ | 0.6 | 0.9 | 0.1-1 |
| $k$ | lysis rate for half-maximal CAR T cell proliferation | cells$^2$/day$^2$ | 2.05e7 | 2.05e7 | |
| $q$ | CAR T cell exhaustion rate | cells$^{-1}$day$^{-1}$ | 1.6e-9 | 1.6e-9 | |
| $m$ | CAR T cell death rate | day$^{-1}$ | 0.293 | 0.293 | |

Table S2: Estimated model parameters to fit model output to murine study data. The range of values allowed for a given parameter in the initial run of the iterative search algorithm is included in the final column. For parameters that were fixed, rather than estimated, the final column is blank.
